# Coordination of -1 Programmed Ribosomal Frameshifting by Transcript and Nascent Chain Features Revealed by Deep Mutational Scanning

**DOI:** 10.1101/2021.03.11.435011

**Authors:** Patrick J. Carmody, Matthew H. Zimmer, Charles P. Kuntz, Haley R. Harrington, Kate E. Duckworth, Wesley D. Penn, Suchetana Mukhopadhyay, Thomas F. Miller, Jonathan P. Schlebach

## Abstract

Programmed ribosomal frameshifting (PRF) is a translational recoding mechanism that enables the synthesis of multiple polypeptides from a single transcript. In the alphavirus structural polyprotein, -1PRF is coordinated by a “slippery” sequence in the transcript, an RNA stem-loop, and a conformational transition in the nascent polypeptide chain. To characterize each of these effectors, we measured the effects of 4,530 mutations on -1PRF by deep mutational scanning. While most mutations within the slip-site and stem-loop disrupt -1PRF, mutagenic effects upstream of the slip-site are far more variable. Molecular dynamics simulations of polyprotein biogenesis suggest many of these mutations alter stimulatory forces on the nascent chain through their effects on translocon-mediated cotranslational folding. Finally, we provide evidence suggesting the coupling between cotranslational folding and -1PRF depends on the translation kinetics upstream of the slip-site. These findings demonstrate how -1PRF is coordinated by features within both the transcript and nascent chain.

## Introduction

Programmed ribosomal frameshifting (PRF) is a translational recoding mechanism that occurs in all kingdoms of life. Though a handful of prokaryotic and eukaryotic PRF motifs have been identified, most of the well-characterized motifs are found within viral genomes (Atkins et al., 2016). Viruses utilize ribosomal frameshifting to increase their genomic coding capacity and to regulate the stoichiometric ratios of viral protein synthesis. Some rely on these motifs to coordinate genomic replication, while others utilize PRF to regulate the production of the structural proteins that mediate assembly (Penn et al., 2020a). In some cases, the frameshift products are themselves virulence factors that antagonize the host interferon response (Hallengard et al., 2014; Rogers et al., 2020; Snyder et al., 2013; Taylor et al., 2016). For these reasons, the efficiency of PRF, which is globally regulated by both host and viral proteins (Napthine et al., 2017; Wang et al., 2019), is often critical for infection and immunity.

The most common type of PRF involves a −1 shift in reading frame (-1PRF) and minimally requires a “slippery” heptanucleotide site in the transcript that typically takes the form X_1_ XXY_4_ YYZ_7_, where X represents three identical nucleotides, Y is an A or U, and Z can be an A, C, or U (Jacks et al., 1988). This structure reduces the thermodynamic penalty associated with non-native codon-anticodon interactions in the −1 reading frame (Bock et al., 2019), which increases the intrinsic efficiency of ribosomal frameshifting by several orders of magnitude (Reil et al., 1993; Stahl et al., 2002). Most - 1PRF motifs also feature an mRNA stem-loop or pseudoknot that forms 6-8 bases downstream of the slip-site (Giedroc and Cornish, 2009; Mouzakis et al., 2013). This structure imposes a mechanical resistance that stalls the ribosome on the slip-site and provides time for the anticodon loops of the A- and/ or P-site tRNA to sample alternative base pairing interactions (Caliskan et al., 2014; Chen et al., 2014; Choi et al., 2020; Kim et al., 2014). Frameshifting is also enhanced by the depletion of the t-RNA pool that decodes the YYZ_7_ codon (“hungry” frameshifting), which provides the opportunity for the P-site tRNA to move out of frame (Korniy et al., 2019b). In addition to mRNA and tRNA effectors, our group recently found that -1PRF is enhanced by mechanical forces generated by the cotranslational folding of the nascent polypeptide chain (Harrington et al., 2020). This finding has since been echoed by recent evidence suggesting cotranslational folding of nsp10 alters -1PRF during translation of SARS-CoV-2 genome (Bhatt et al., 2020). The efficiency of - 1PRF can also be modified in *trans* by a variety of host and viral proteins and nucleic acids (Penn et al., 2020a). Thus, the efficiency of ribosomal frameshifting can be modified by a wide variety of effectors that tune the rate of translation, the structural properties of the transcript, and/ or the conformation of the nascent chain. Nevertheless, a unified understanding of how these features cooperate to establish the efficiency of PRF is needed (Mikl et al., 2020).

In this work, we utilize deep mutational scanning (DMS) to identify sequence constraints associated with -1PRF in the multipass integral membrane protein known as the Sindbis virus (SINV) structural polyprotein. We report the effects of 4,530 mutations on the relative efficiency of -1PRF in the context an extended genetic reporter containing the native features that coordinate the biosynthesis and cotranslational folding of the nascent polyprotein at the endoplasmic reticulum (ER) membrane (Fig. 1A). Incorporating these elements into our reporter facilitated the identification of -1PRF modulators in both the transcript and within several regions of the nascent chain. As expected, mutational patterns within the slip-site and the region immediately downstream of the slip-site are consistent with the formation of a previously characterized RNA stem-loop (Chung et al., 2010; Kutchko et al., 2018). However, we find that -1PRF is also sensitive to numerous mutations within the ~160 base region preceding the slip-site, which encode a pair of transmembrane (TM) domains that occupy the ribosomal exit tunnel and translocon during frameshifting. In conjunction with a suite of atomistic and coarse-grained molecular dynamics (CGMD) simulations (Niesen et al., 2020), we show that many mutations within this region attenuate -1PRF through their effects on a folding intermediate that forms during translocon-mediated membrane integration of the nascent polyprotein. We also find that -1PRF appears to be sensitive to changes in the translation kinetics within the region immediately preceding the slip-site, which influences the pulling forces generated by the cotranslational folding of the nascent chain. Together, these findings provide a detailed description of the sequence constraints of -1PRF within the alphavirus structural polyprotein.

**Figure 1.**
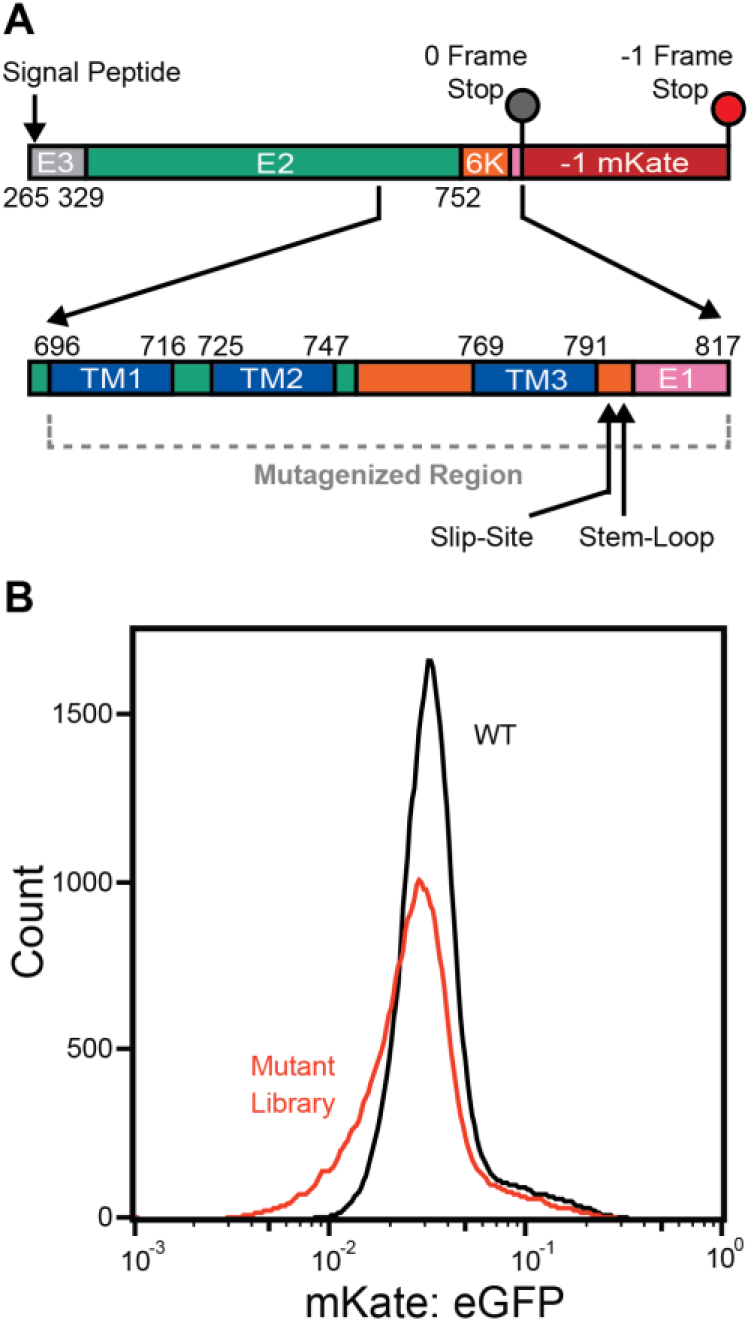
Library of ribosomal frameshifting reporter variants. A) A cartoon depicts the structure of a reporter that generates a fluorescent protein (mKate) as a result of -1PRF in the SINV structural polyprotein (top). A library of reporter variants was generated by randomizing every 0-frame codon within a region ranging from residues 696-817 (bottom). B) A histogram depicts the range of single-cell mKate: GFP intensity ratios among recombinant stable cells expressing the WT reporter (black) or the library of reporter variants (red).

## Results

### Production of a stable HEK293T cell line expressing a programmed array of -1PRF reporter variants

To probe the sequence constraints of -1PRF, we employed DMS to map the effects of mutations on ribosomal frameshifting within the SINV structural polyprotein. Mutagenic effects were assessed in the context of a reporter that selectively generates a fluorescent protein (mKate) as a result of -1PRF during translation and processing of the SINV structural polyprotein (Fig. 1A). The design of this reporter is similar, in principle, to those described in a recent high-throughput investigation of PRF (Mikl et al., 2020), except that we included a much larger fragment of the transcript (1.7 kb, 557 amino acids) and opted to track expression using an IRES-eGFP cassette rather than an N-terminal fusion domain. This design preserves the integrity of the native signal peptide in the E3 protein (Fig. 1A), which is critical for the targeting and maturation of the nascent polyprotein at the ER membrane. To survey the effects of mutations on the reporter signal, we first generated a one-pot genetic library by randomizing each 0-frame codon within a region beginning at the N-terminal codon of the first TM domain (TM1) of the E2 protein (I696) and ending 23 codons downstream of the slip-site (P817, Fig. 1A). We then used this genetic library (7,614 total variants) to generate recombinant stable HEK293T cell lines in which each cell expresses a single variant from a unique Tet-inducible promoter within the genome, as previously described (Matreyek et al., 2017). Importantly, the specificity of this recombination reaction ensures individual variants are expressed at similar levels within each cell. Nevertheless, we normalized mKate levels according to the IRES-eGFP intensity to account for small variations in expression, as has been previously described (Chiasson et al., 2020).

On average, stable cells expressing individual single-codon variants have an mKate: eGFP intensity ratio that is comparable to that of cells expressing the WT reporter (Fig. 1B). However, a sub-set of these cells express variants that generate intensity ratios outside the WT intensity range (Fig. 1B). This suggests that, while most mutations have minimal impact, the library contains many that modify the efficiency of -1PRF. To analyze the effects of individual mutations, we fractionated these cells according to their mKate: eGFP intensity ratios using fluorescence activated cell sorting (FACS), quantified the variants within each fraction by deep sequencing, then used these data to estimate the intensity ratio for each individual variant as described previously (Penn et al., 2020b). Intensity ratios were generally consistent across biological replicates when normalized according to the WT value (Pearson’s R = 0.79-0.83, Fig. S1A-C). Intensity ratios determined by DMS were also correlated with independent measurements in cells transiently expressing individual reporter variants (Fig. S1D). Together, our results reveal how mutations upstream of the slip-site, downstream of the slip-site, and within the slip-site itself impact ribosomal frameshifting (Fig. 1A).

### Sequence constraints within the heptanucleotide slip-site

Slippery nucleotide sequences enable favorable codon-anticodon base pairing in the −1 reading frame, and the efficiency of -1PRF is therefore highly sensitive to mutations that disrupt the slip-site (Brierley et al., 1992). As expected, we find that every single-nucleotide variant (SNV) within the heptanucleotide slip-site (U_1_ UUU_4_ UUA_7_) measurably decreases -1PRF (Fig. 2A, yellow). Furthermore, an analysis of 128 single, double, and triple mutants within the slip-site reveals that frameshifting decreases with an increasing mutational load within the UUU_4_ (P-site) or UUA7 (A-site) codons (Fig. 2B). These mutations presumably decrease the efficiency of -1PRF by increasing the free energy difference between the codon-anticodon base pairing in the 0 and −1 reading frames. However, we also find frameshifting to be sensitive to mutations within the adjacent codon upstream of the slip-site (Fig. 2A). This observation potentially suggests the observed PRF signal may arise from some combination of −1 and −4 frameshifting transitions, as has been observed for other PRF motifs (Yan et al., 2015). It is also possible that the observed - 1PRF signal arises from a mix of canonical frameshifting (two-tRNA) and “hungry” frameshifting (one-tRNA), where the P-site tRNA frameshifts prior to the decoding of the UUA7 codon. The overall efficiency of both the HIV and Semliki forest virus -1PRF motifs is maximized when the relative abundance of the tRNA that decodes the UUA7 codon is depleted (Korniy et al., 2019b). Similarly, we find that -1PRF is decreased by all three A7 mutations in the SINV slip-site (Fig. 2A), which should attenuate hungry frameshifting given that these codons are decoded by tRNA that are more abundant in these cells (Kirchner et al., 2017; Korniy et al., 2019a). Together, these observations highlight the essential role of the slip-site and its adjacent bases, yet suggest the observed -1PRF signal potentially arises from a spectrum of frameshifting transitions.

**Figure 2.**
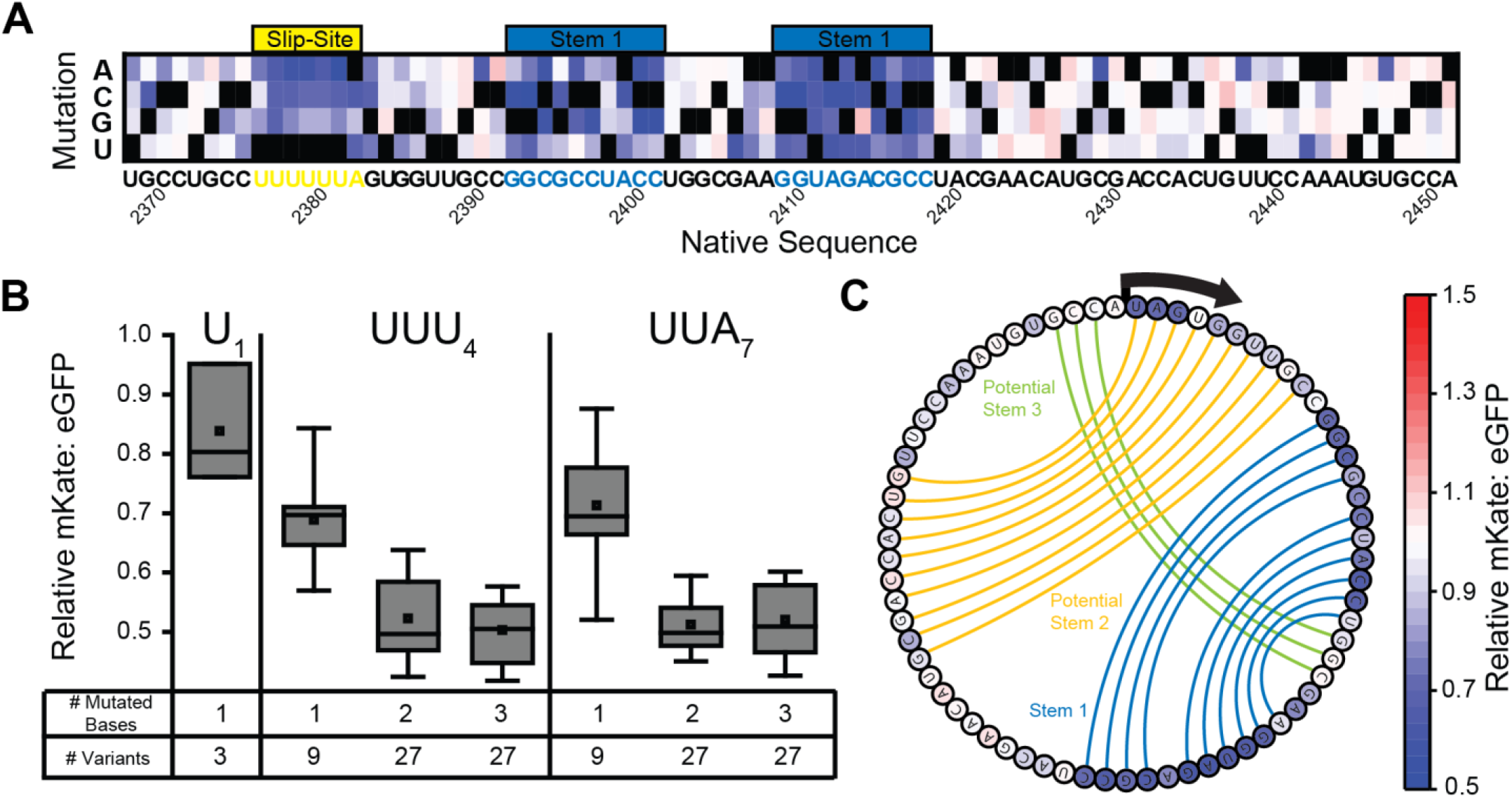
Impact of slip-site and stem-loop mutations on ribosomal frameshifting. A) A heatmap depicts the effects of every single nucleotide substitution (Y-coordinate) at each position (X-coordinate) within or near the slip-site on the mKate: eGFP ratio as determined by DMS. Ratios were normalized relative to that of WT, and black squares are shown in place of the native nucleobase at each position. The relative position of the slip-site (yellow) and stem 1 (blue) are shown for reference. B) A box plot depicts the distribution of relative mKate: eGFP values associated with variants bearing one, two, or three mutations at certain positions within the slip-site. The upper and lower boundaries of the box correspond to the 75^th^ and 25^th^ percentile values, and the upper and lower whiskers correspond to the 90^th^ and 10^th^ percentile values, respectively. The horizontal line within the box corresponds to the median value, and the square corresponds to the position of the average value. C) Connecting lines depict previously predicted base pairing interactions within stem 1 (blue), potential stem 2 (orange), and potential stem 3 (green) are shown in the context of the nucleotide sequence downstream from the slip-site (Chung et at., 2010). Each nucleotide is colored according to the average relative mKate: eGFP intensity ratio across the three mutations at each position.

### Sequence constraints within the stimulatory RNA stem-loop

Efficient ribosomal frameshifting requires a stimulatory RNA secondary structure that impedes translocation and increases the dwell time of the ribosome on the slip-site. Stimulatory structures within the sub-genomic RNA of a number of alphaviruses have been previously characterized (Chung et al., 2010; Kendra et al., 2018; Kutchko et al., 2018). Secondary structure predictions for SINV suggest the region adjacent to the slip-site is capable of forming a pseudoknot consisting of one primary stem-loop (stem 1), a secondary stem-loop within the spacer region (potential stem 2), and a short stem that basepairs within the loop of stem 1 (potential stem 3) (Chung et al., 2010). However, experimental evidence suggests frameshifting is primarily driven by stem 1 (Chung et al., 2010). Consistent with these findings, our map of mutational effects shows that frameshifting is highly sensitive to mutations within a region beginning 8 bases downstream of the slip-site that corresponds to the position of stem 1 (Figs. 2 A & C). Furthermore, the contacts suggested by this map correspond exactly to those identified by SHAPE analysis (Kutchko et al., 2018). Interestingly, stem 1 contains a single mismatch (Fig. 2C), and PRF is enhanced by the only SNV that creates a G-C base pair at this position (2414 A→G, Fig. 2A). We also find that frameshifting is enhanced by two SNVs that extend the base of the stem-loop by an additional base pair (2391 C→A, 2419 U→G, Fig. 2A). Together, these observations confirm that stem 1 is critical for PRF efficiency. While more stable equilibrium structure(s) could potentially form, our functional analysis suggests stem 1 is the dominant structure within the ensemble that stimulates -1PRF between rounds of ribosome-mediated unwinding (Halma et al., 2019; Ritchie et al., 2012).

### Sequence constraints within the nascent polypeptide chain

Our recent investigations of the impact of the nascent chain on -1PRF revealed that the mechanical force generated by the translocon-mediated membrane integration of the second TM domain within the E2 protein enhances ribosomal frameshifting in the SINV structural polyprotein (Harrington et al., 2020). Our DMS results reveal that ribosomal frameshifting is sensitive to a variety of mutations that alter the amino acid sequence within the portion of the nascent chain that has been translated at the point of frameshifting, including mutations to residues within the exit tunnel, between the exit tunnel and translocon, and within the second transmembrane domain of E2 (TM2, Fig. 3). Consistent with previous findings (Harrington et al., 2020), our results show that the introduction of polar or charged side chains within a certain region of TM2 reduces -1PRF (V735-V746) while mutations that enhance the hydrophobicity of TM2 generally increase -1PRF (Fig. 3). However, the effects of mutations on -1PRF appear to deviate from their predicted effects on the energetics of translocon-mediated membrane integration (Fig. S2). Mutations to residues near the center of TM2 are predicted elicit the largest changes in the energetics of translocon-mediated membrane integration (A734 - T738, Fig. 4A, red), which reflects the depth-dependence of amino acid transfer free energies (Hessa et al., 2007; Moon and Fleming, 2011). In contrast, variations in mKate: eGFP intensity ratios are most pronounced among variants bearing mutations within the C-terminal residues of TM2 (Fig. 4A, black). This discrepancy suggests that, in addition to the hydrophobicity of TM2 and its corresponding transfer free energy, stimulatory pulling forces may hinge upon structural and/ or dynamic constraints associated with its translocon-mediated membrane integration.

**Figure 4.**
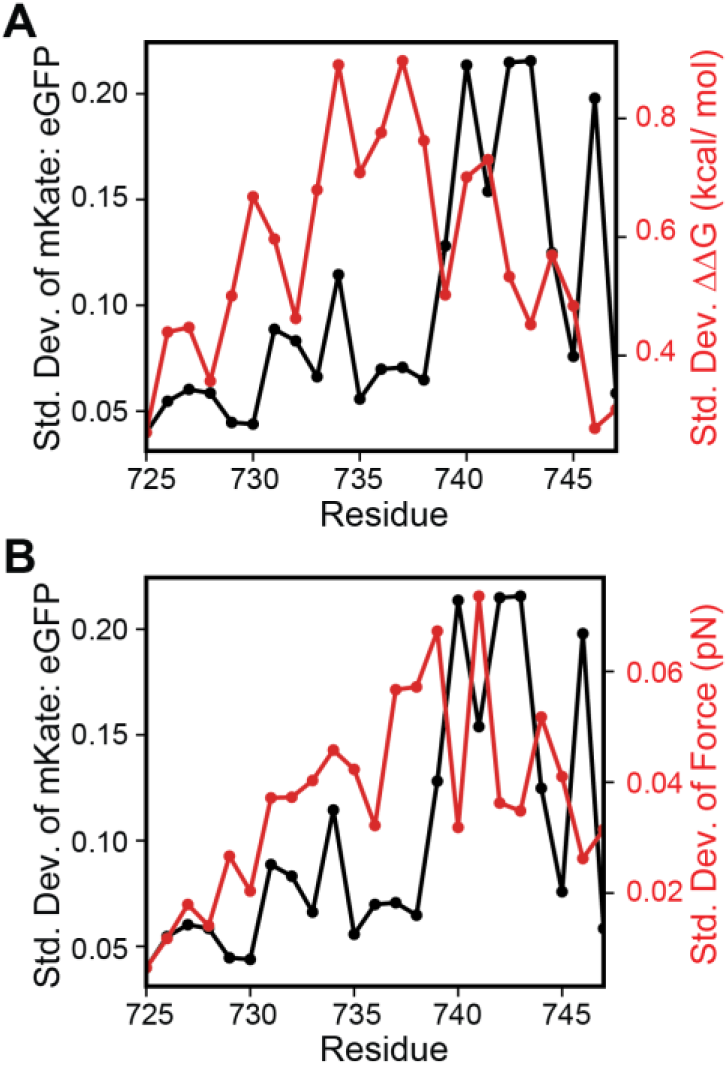
Impact of TM2 mutations on -1PRF, membrane integration energetics, and pulling forces. The standard deviation of the relative mKate: eGFP intensity ratios for all 19 amino acids substitutions (black) at each residue are plotted in relation to the corresponding standard deviation of (A) the predicted change in the transfer free energy associated with the translocon-mediated membrane integration of TM2 as determined by the ΔG predictor (red) (Hessa et al., 2007) and B) the relative pulling force generated by its translocon-mediated membrane integration as was determined by CGMD simulations (red).

**Figure 3.**
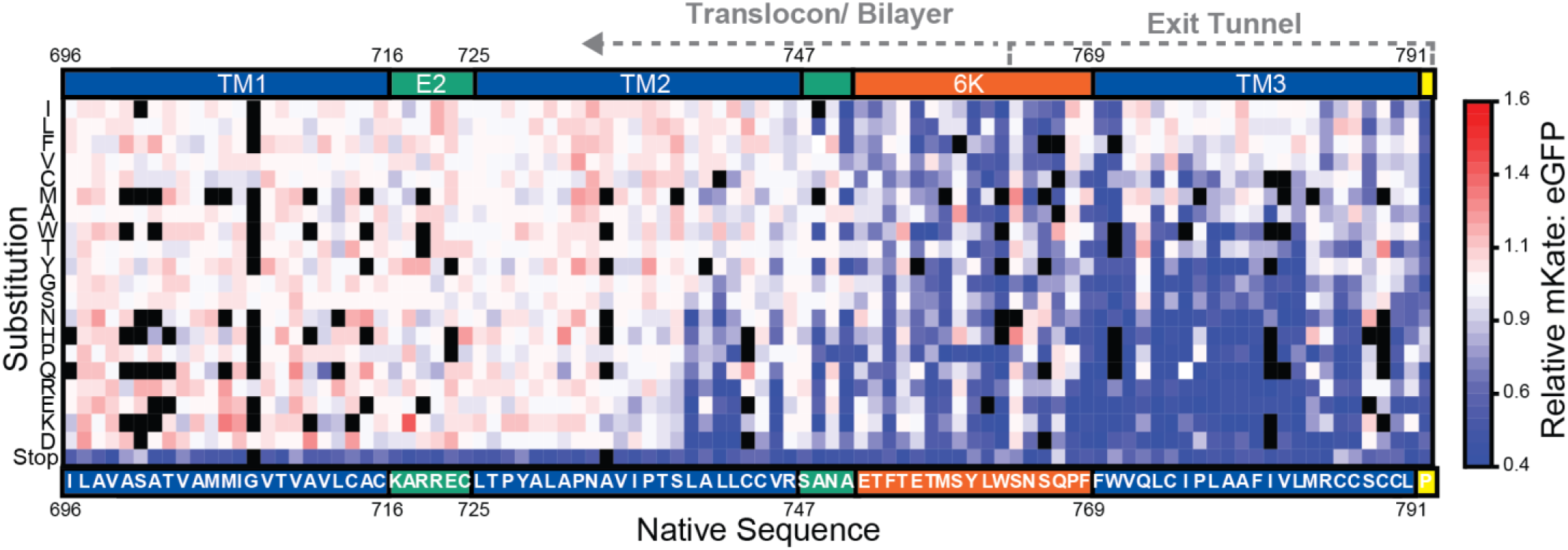
Impact of nascent chain mutations on ribosomal frameshifting. A heatmap depicts the effects of every amino acid substitution (Y-coordinate) at each position (X-coordinate) within the nascent polypeptide chain on the mKate: eGFP ratio as determined by DMS. Substitutions are arranged from most hydrophobic (top) to most polar (bottom). Ratios were normalized relative to that of WT, and black squares indicate mutations that lack coverage. The positions of each TM domain (blue) and the slip-site (yellow) as well as the domain boundaries for the E2 (green) and 6K (orange) proteins are shown for reference. The amino acid that corresponds to the first codon of the slip site (containing U1) is indicated in yellow. The residues that are likely to reside within the exit tunnel or outside of the ribosome during ribosomal frameshifting are indicated with dashed lines.

To explore the effects of mutations on nascent chain pulling forces, we performed CGMD simulations of polyprotein translation that include coarse-grained representations of the ribosomal exit tunnel and the Sec translocon in an implicit lipid bilayer (Niesen et al., 2017). The nascent polypeptide is treated as a polymer of beads (3 AA/ bead) that emerge from the ribosome exit tunnel at the physiological rate of translation (5 AA/ s). The hydrophobicity and charge of each bead is derived from its three constituent amino acids. To quantify stimulatory pulling forces, we paused translation for three seconds once the ribosome reaches the slip-site and measured the force on the nascent chain as it explores the environment in, on, and around the translocon as has been previously described (Niesen et al., 2018). Consistent with previous findings (Harrington et al., 2020), simulations of the 437 missense variants bearing mutations within TM2 reveal that mutations that attenuate pulling forces on the nascent chain generally reduce frameshifting; pulling forces are correlated with experimentally derived relative mKate: eGFP intensity ratios (Pearson’s R = 0.48, Fig. S3A). Hydrophobic substitutions in TM2 increase the propensity of TM2 to enter the translocon and partition into the membrane in these simulations, which generates higher pulling forces (Fig. S3B) and enhanced frameshifting (Fig. S3C). Conversely, polar mutations decrease translocon occupancy, membrane integration of TM2, and pulling forces. Importantly, both pulling forces from CGMD and the observed mKate: eGFP intensity ratios are most sensitive to mutations within the C-terminal residues of TM2 (Fig. 4B, red). This agreement suggests stimulatory forces originate from conformational transitions that are mediated by dynamic interactions between TM2, the translocon, and the lipid bilayer.

To evaluate the structural context of mutations that influence -1PRF, we constructed an atomistic model of TM2 nested within the translocon. An initial model was generated by mapping the polyprotein sequence onto a cryo-EM structure of a nascent chain intermediate that adopts an Nin topology comparable to that of TM2 in the context of the translocon (Ma et al., 2019). After a 150 ns equilibration of the translocon-nascent chain complex within an explicit lipid bilayer, TM2 adopts a tilted orientation in which the N-terminal portion of TM2 projects into the lipids while its C-terminal residues remain wedged within the lateral gate of the translocon (Fig. 5A). The N-terminal residues that interact with the bilayer are generally more dynamic than the C-terminal residues that interact with the translocon (Fig. 5B). TM2 achieves a similar topological orientation and exhibits similar dynamic fluctuations regardless of how the amino acid sequence is initially mapped onto the structural template of the nascent chain (Fig. S4). While the magnitude of the dynamic fluctuations observed on the time scales of the atomistic simulations is modest relative to coarse-grained simulations, the conformational dynamics observed within both models indicate that fluctuations within the C-terminal residues of TM2 are suppressed by interactions with the translocon (Fig. 5C), which is the final portion of the helix to partition into the bilayer. Mutations that enhance the polarity of these residues generally decrease -1PRF (Fig. 5A), presumably by strengthening interactions with the translocon and/ or disfavoring the transfer of these residues into the bilayer. Taken together, these observations suggest mutations in TM2 influence pulling forces, and by extension -1PRF, by modifying interactions that form between the translocon and nascent chain.

**Figure 5.**
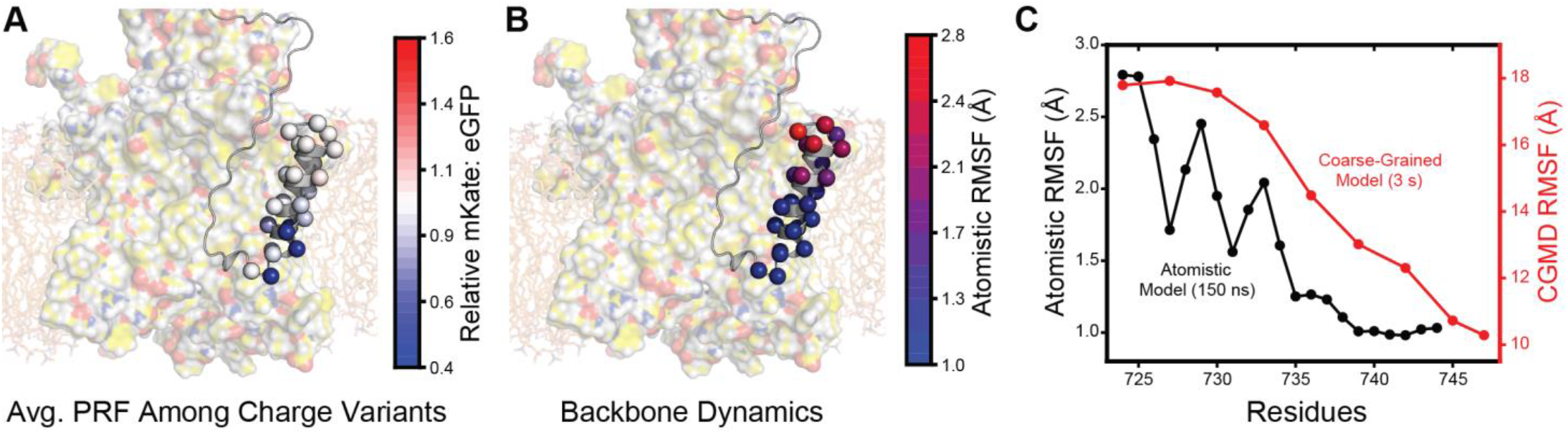
Structural context and conformational dynamics of nascent TM2. **A)** The average relative mKate: eGFP intensity ratios associated with mutations that introduce charged side chains at each position within TM2 are mapped on to an atomistic model of nascent TM2 at the approximate point of frameshifting. Residues are rendered as spheres and colored based on the relative mKate: eGFP ratio for charged substitutions as was determined by DMS. The image of the nascent chain structure is superimposed on top of the structure of the translocon and lipid bilayer for clarity. B) The root mean square fluctuation (RMSF) of each Cα during a 150 ns all-atom molecular dynamics simulation are mapped on to an atomistic model of nascent TM2 at the approximate point of frameshifting. Residues are rendered as spheres and colored according to RMSF. The image of the nascent chain structure is superimposed on top of the structure of the translocon and lipid bilayer for clarity. C) RF values from atomistic (Cα, black) and coarse-grained (bead indexed to sequence, red) molecular dynamics simulations are plotted for each position within TM2.

### Impact of translation kinetics on pulling forces and -1PRF

Ribosomal frameshifting is also highly sensitive to mutations within the region of the transcript encoding the residues between TM2 and the peptidyl transfer center (S748 – L791, Fig. 3). Most of these mutations alter the portion of the nascent chain that resides within the ribosomal exit tunnel during - 1PRF. With respect to the amino acid sequence, these mutagenic trends suggest native -1PRF levels are only maintained by mutations that preserve the hydrophobicity of this segment (Fig. 3). This may suggest this segment is capable of forming structural contacts within the exit tunnel (see *Discussion*). Nevertheless, it should also be noted that the translation of this segment occurs as TM2 begins to emerge from the exit tunnel and interact with the translocon and/ or membrane. Modifications that alter the translation kinetics within this region could therefore also impact the membrane integration of TM2 and pulling forces on the nascent chain (Niesen et al., 2020). To evaluate the potential role of translational kinetics, we calculated mKate: eGFP intensity ratios associated with each individual codon substitution, then sorted the values for each codon according to the relative abundance of their decoding tRNAs in HEK293 cells (see *Methods*). A heat map of intensity ratios reveals that -1PRF is generally attenuated by mutations within the 21 codons upstream of the slip-site that increase the relative abundance of decoding t-RNA (Fig. 6). Codons that are decoded by low-abundance tRNAs are generally tolerated within this region (Fig. 6), which suggests that efficient -1PRF hinges upon the slow translation of these codons.

**Figure 6.**
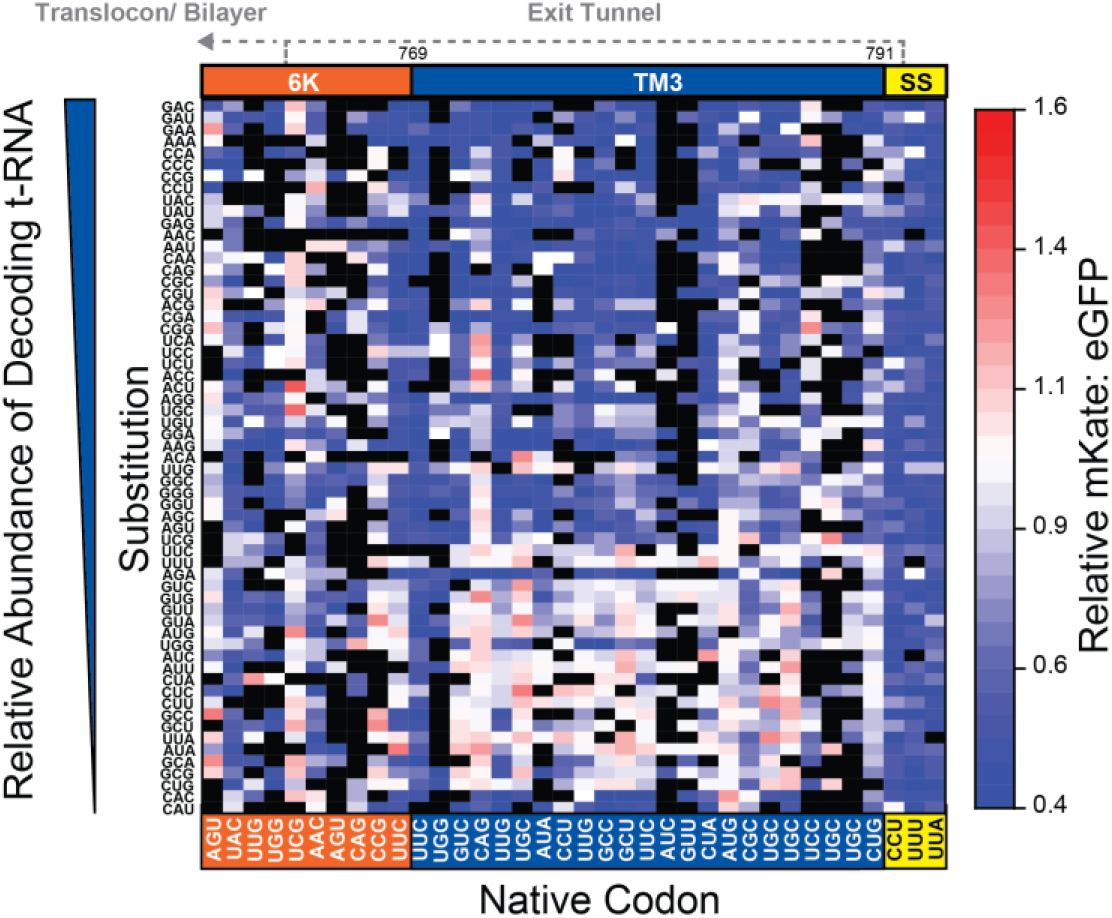
Impact of codon modifications on ribosomal frameshifting. A heatmap depicts the effects of every codon substitution (Y-coordinate) at each 0-frame codon (X-coordinate) within the nascent polypeptide chain on the mKate: eGFP intensity ratio as determined by DMS. Substitutions are arranged with respect to the relative abundance of tRNA that can decode each codon in HEK293 cells (see *Methods*). Ratios were normalized relative to that of WT, and black squares indicate mutations that lack sufficient coverage. The positions of TM3 (blue), the preceeding 6K residues (orange), and the slip-site (yellow) are shown for reference. The residues that are likely to reside within the exit tunnel or outside of the ribosome during ribosomal frameshifting are indicated with dashed lines.

Mutations that increase the relative abundance of decoding tRNAs and accelerate translation could potentially decrease - 1PRF by reducing the efficiency of translocon-mediated membrane integration (Niesen et al., 2020). To assess how translational kinetics impact the membrane integration of TM2, we carried out a series of CGMD simulations of polyprotein biosynthesis in which the rate of translation was varied. An analysis of these trajectories reveals that the rate of translation has minimal impact on the final topology of TM2 (Fig. S5). Nevertheless, -1PRF is likely to be more closely associated with the magnitude of the pulling forces on the nascent chain rather than the equilibrium structure of TM2. To evaluate whether pulling forces are sensitive to translation kinetics, we varied the rate of translation within CGMD simulations and measured the pulling force on the nascent chain while the ribosome occupies the slip-site. One hundred independent trajectories were carried out at distinct translation rates ranging from 1 to 15 amino acids per second, and pulling forces were averaged over 0.8 s once the ribosome reaches the slip-site. A plot of the mean force against translation rate reveals that the pulling force trends upward as translation slows. These results provide secondary evidence suggesting the mechanical forces that stimulate -1PRF are likely attenuated at higher translation rates. In conjunction the observed mutagenic trends (Fig. 6), our findings suggest that efficient frameshifting may be more efficient when translation of the region preceding the slip-site is slow.

## Discussion

The interactions between the transcript and translation machinery that stimulate -1PRF have been studied in considerable detail (Atkins et al., 2016; Korniy et al., 2019b). However, our recent investigations of the alphavirus structural polyprotein revealed that -1PRF is also sensitive to conformational transitions in the nascent chain (Harrington et al., 2020). The net efficiency of -1PRF in this system is therefore modulated by the interplay between structural features within both the transcript and nascent chain. To map these effectors, we measured the effects of 4,530 mutations on the efficiency of -1PRF in the context of a reporter containing a large fragment of the SINV structural polyprotein. This reporter retains all the native features that direct the targeting, processing, and cotranslational folding of the nascent chain (including the signal peptide and post-translational modification sites) as well as the features in the transcript that stimulate -1PRF (slip-site and stem-loop). Retaining these features allowed us to identify and map effectors in a manner that cannot be achieved using a conventional dual-luciferase reporter system bearing an N-terminal fusion domain (Fig. S6). Our observed trends validate numerous expectations about the stimulatory RNA elements. For instance, all 128 single, double, and triple mutants that alter the heptanucleotide slip-site reduce - 1PRF to some extent (Fig. 2 A & B). Additionally, the mutagenic trends downstream from the slip-site are entirely consistent with previous characterizations of the stimulatory stem-loop (Fig. 2C) (Chung et al., 2010; Kutchko et al., 2018). Interestingly, the latter measurements also show that -1PRF can be enhanced by mutations that strengthen or extend this stem-loop (Fig. 2A). The fact that the stability of this stem-loop has not evolved to maximize -1PRF provides a clear example of how SINV has evolved to maintain a specific frameshifting efficiency.

We recently showed that ribosomal frameshifting in the SINV structural polyprotein is sensitive to mutations that perturb the translocon-mediated membrane integration of TM2 (Harrington et al., 2020). Considering the marginal hydrophobicity of this segment, we initially expected mutagenic trends within this region to reveal a simple relationship between the topological energetics of TM2 and the efficiency of -1PRF. However, the observed mutagenic trends cannot be explained by hydrophobicity alone (Fig. S2). Instead, our DMS and molecular dynamics data suggest that stimulatory pulling forces are specifically generated by the transfer of the C-terminal portion of TM2 from the translocon to membrane. At this final stage of translocation, these residues interact with the lumenal edge of the lateral gate, which pulls the loop between TMs 2 & 3 into the protein conducting channel of the translocon (Fig. 5). This loop is wedged between TM2 and the translocon at the point of frameshifting, and the observed mutagenic patterns within this region do not simply track with hydrophobicity or codon usage (Fig. 3 & 6). This observation suggests this segment may form specific interactions that influence membrane integration. Nevertheless, it is unclear whether mutations within this loop alter -1PRF by reducing the force on the nascent chain or by delaying the transmission of the force to the peptidyl transfer center until after the ribosome passes the slip-site. Additional investigations are needed to explore the network of interactions between the nascent chain, translocon, and lipid bilayer that mediate this type of mechanochemical feedback (Leininger et al., 2019).

The ribosomal exit tunnel generally suppresses protein structure formation until the nascent chain clears the ribosome (Kudva et al., 2018; Samelson et al., 2016). We were therefore surprised to find that -1PRF is highly sensitive to mutations within the region encoding the portion of the nascent chain that occupies the exit tunnel during frameshifting (Fig. 3). Nevertheless, a recent investigation of the sequence constraints in other PRF motifs found that frameshifting is often enhanced by rare codons upstream of the slip-site (Mikl et al., 2020). An analysis of the effects of codon-level substitutions suggests changes in -1PRF may be related to the effects of mutations on the relative abundance of decoding tRNAs (Fig. 6), and CGMD simulations suggest stimulatory forces are sensitive to the rate of translation (Fig. 7). These observations strongly suggest translation kinetics are critical for the coupling between cotranslational folding and -1PRF. However, it remains unclear how much these substitutions actually impact the physiological rate of translation. Furthermore, the estimated impact of translation kinetics on pulling forces appears to be relatively modest in relation to the magnitude of their effects on -1PRF. Given these caveats, it may be that these mutations impact -1PRF in more than one way. For instance, the native codon bias in this region may also control the frequency of ribosomal collisions, which were recently found to stimulate -1PRF (Smith et al., 2019). Modifications to the structure of the nascent chain could also potentially be relevant to -1PRF. This region encodes TM3, and it was recently found that hydrophobic helices exhibit a tendency to form helical structure within the exit tunnel (Bano-Polo et al., 2018). Furthermore, native -1PRF levels are only maintained when mutations introduce other hydrophobic side chains within this segment (Fig. 3). Thus, the formation of helical structure within the exit tunnel could potentially also contribute to this mechanochemical coupling. Taken together, it seems the mechanistic basis of the mutagenic effects within this region are undoubtedly complex, and additional investigations are needed to tease apart each of these variables.

**Figure 7.**
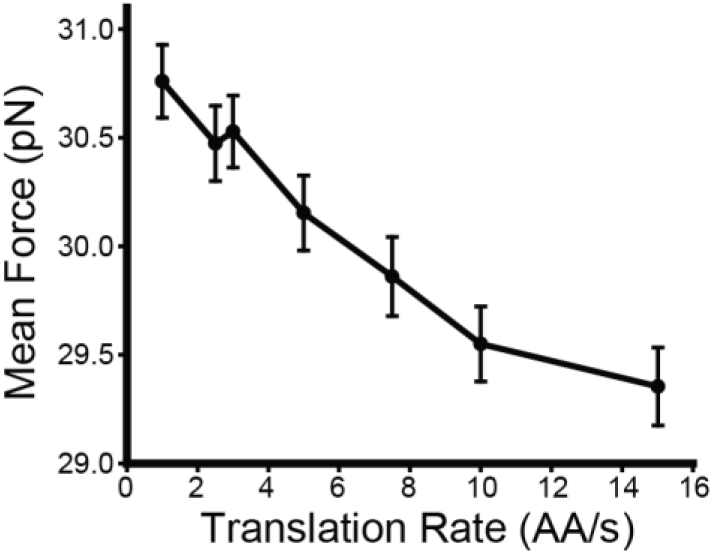
Impact of translation kinetics on nascent chain pulling forces. Coarse-grained molecular dynamics simulations of the translocon-mediated cotranslational folding of the SINV structural polyprotein were carried out varying the translation rate. Pulling forces on the nascent chain were recorded over 0.8 s while the ribosome occupies the slip-site. The mean force on the nascent chain across 100 independent trajectories is plotted against the corresponding translation rate. Error bars reflect the standard error of the mean.

Together, our results reveal that the net efficiency of -1PRF in the alphavirus structural polyprotein arises from the coupling between multiple structural effectors. While this motif features the canonical slip-site and stem loop architecture found in most -1PRF sites, its efficiency is rendered tunable by mutations that alter mechanochemical transitions within the nascent chain and/ or the translation kinetics upstream of the slip-site. Our DMS data demonstrate that this degeneracy effectively expands the pool of genetic variations that are capable of adjusting the efficiency of -1PRF; an important consideration for viral evolution. Though this investigation focuses on a mechanochemical transition within the ribosome-translocon complex, it seems likely that many other classes of conformational transitions within the nascent chain may also be capable of stimulating -1PRF. Further investigations are needed to evaluate whether similar mechanisms are operative within other transcripts.

## Methods

### Plasmid and Library Preparation

-1PRF measurements were made using a previously described fluorescent reporter that selectively generates a fluorescent mKate protein as a result of -1PRF during translation of the SINV 6K gene (Harrington et al., 2020). To control for variations in expression, the intensity of the mKate -1PRF signal is measured in relation to the eGFP intensity generated by a downstream IRES-eGFP cassette. To adapt this reporter for DMS, the CMV promoter and enhancer within this vector were replaced with an attB recombination site using NEBuilder. An Nt.BbvCI/ Nb.BbvCI endonuclease site was also introduced into the backbone using site-directed mutagenesis in order to facilitate the production of one-pot genetic libraries. A library of single-codon variants was generated using nicking mutagenesis (Wrenbeck et al., 2016). Briefly, the sense strand of the parental plasmid was nicked and degraded. The antisense strand was then used as template for the synthesis of a complementary strand using a pool of mutagenic oligonucleotides that each contain a randomized NNN codon. The WT antisense strand of the heteroduplex was nicked and degraded. The remaining mutagenized sense strand library was then used as a template for synthesis using a universal primer that anneals outside of the mutagenized region. The resulting plasmid library was then electroporated into electrocompetent NEB10β cells (New England Biolabs, Waltham, MA), which were then grown in liquid culture prior to purification of the plasmid library using a ZymoPure II Endotoxin-Free Midiprep kit (Zymo Research, Irvine, CA). An analysis of our final DMS data suggest this library contains at least 7,614 of 7,686 possible variants. Nevertheless, in this manuscript we have chosen to focus on the effects of 4,530 of these mutations within specific regions of interest, which are shown in the heatmaps in Figures 2A, 3, and 6. To validate DMS measurements, a representative series of individual variants were introduced into the originally described expression vector using site-directed mutagenesis (Harrington et al., 2020). To compare -1PRF levels in the context of a dual luciferase reporter, the fragment of the SINV structural polyprotein shown in Fig. 1A (residues 265-817) was inserted into a dual-luciferase reporter construct (pJD2257) (Kendra et al., 2018) that was modified in order to remove a cryptic splice site in the Renilla luciferase gene (Khan, 2019).

### Cell Culture and Production of Recombinant Stable Cell Lines

All cells were grown in Dulbecco’s modified eagle medium (Gibco, Carlsbad, CA) supplemented with 10% fetal calf serum (Corning, Corning, NY), and penicillin (100 U/ml)/ streptomycin (100 μg/ml) at 37°C in a humidified incubator containing 5% CO2 by volume. Recombinant stable cell libraries expressing a mix of polyprotein reporter variants from a uniform genomic locus were generated by cotransfecting a genetically-modified HEK293T cell line with the plasmid library and a Bxb1 expression vector as was described previously (Matreyek et al., 2017; Penn et al., 2020b). Cells that underwent the desired recombination reaction were isolated using a FACS Aria II cell sorter (BD Biosciences, Franklin Lakes, NJ) based on the characteristic gain in eGFP and loss of blue fluorescent protein (BFP) expression that occurs as a result of the insertion a single copy of the reporter downstream from the Tet-inducible promoter within the genomic “landing pad” (Matreyek et al., 2017). Following isolation, recombinant cells were grown in media containing 2 μg/ mL doxycycline in order to maintain the stable expression of polyprotein variants from the genomic landing pad.

### Deep Mutational Scanning

A minimum of 2 million stably recombined cells were isolated by FACS to ensure the library of stable cells represents an exhaustive sampling of the variants within the plasmid library (~1,000 X coverage). This population was then expanded prior to its fractionation into quartiles (~2 million cells per fraction) according to the mKate: eGFP intensity ratio by FACS. Each cellular fraction was then expanded and harvested prior to the extraction of genomic DNA using the GenElute Mammalian Genomic DNA Miniprep kit (Sigma-Aldrich, St. Louis, MO). A semi-nested PCR approach was then used to selectively amplify the mutagenized region of the polyprotein reporter from the genomic integration site in a similar manner as was previously described (Matreyek et al., 2017; Penn et al., 2020b). Amplicons containing the mutagenized regions were prepared from each cellular isolate and sequenced using paired end reads on a 600 cycle Illumina MiSeq cell (Illumina Inc., San Diego, CA) to a final depth of ~2 million reads per amplicon/ isolate, which were filtered based on quality score. The mKate: eGFP intensity ratio for each variant was then inferred from the number of sequence-based identifications within each cellular isolate and normalized relative to the value of WT, as was previously described (Penn et al., 2020b). It should be noted that, due to the nature of the scoring system, the lowest possible score for mKate: eGFP intensity ratios is 0.35, which corresponds to the average intensity ratio of the lowest FACS bin divided by the intensity ratio of WT from each experiment (Penn et al., 2020b).

Relative mKate: eGFP intensity ratios for codon substitutions were mapped according to the relative abundance of decoding tRNAs (Fig. 6) based on previous measurements of the relative abundance of tRNAs in HEK293 cells (Kirchner et al., 2017). Briefly, we averaged previously reported replicate measurements of the relative abundance of tRNAs in HEK293 cells. These values were then used to calculate a score based on the sum of the abundances for all tRNAs that are capable of decoding each codon.

### -1PRF Reporter Measurements

In order to validate DMS measurements, we characterized a series of individual variants using both a fluorescence-based reporter and a dual-luciferase reporter for -1PRF in the SINV 6K gene. Fluorescence reporters bearing polyprotein variants of interest were transiently expressed in HEK293T cells, and the relative mKate: eGFP intensity ratios were determined by flow cytometry as previously described (Harrington et al., 2020). HEK293T cells were transiently transfected with dual-luciferase reporter constructs using lipofectamine 3,000 (Invitrogen, Carslbad, CA) in accordance with the manufacturer’s instructions. Cells were harvested two days post-transfection and lysed using the Passive Lysis Buffer provided in the Dual-Luciferase Reporter Assay System (Promega, Madison, WI). The relative activities of the firefly luciferase and renilla luciferase domains in the clarified lysate were then measured using the Dual-Luciferase Reporter Assay System on a Synergy Neo2 plate reader (Biotek, Winooski, VT) in accordance with the manufacturer’s instructions. To determine the -1PRF efficiency, the firefly: renilla luciferase ratio was normalized relative to that of a construct lacking the insert as was previously described (Jacobs and Dinman, 2004).

### Computational Predictions of Topological Energetics

The effects of mutations on the free energy associated with the transfer of TM2 from the translocon to the ER membrane were estimated using the ΔG predictor (Hessa et al., 2007) using a series of individual predictions for TM2 variants assuming a WT sequence of LTPYALAPNAVIPTSLALLCCVR.

### Coarse-grained simulations of polyprotein translation

Coarse-grained simulations use a previously developed and validated methodology (Harrington et al., 2020; Niesen et al., 2017; Niesen et al., 2018). Simulations are based on coarsened representations of the ribosome exit tunnel, Sec translocon, and nascent chain. The nascent chain is represented as a polymer of beads, each of which represents three amino acids. Each bead has a hydrophobicity and charge derived from its three constituent amino acids. The solvent and lipid bilayer are modelled implicitly. The lateral gate of the Sec translocon stochastically switches between open and closed conformations with probability dependent on the free energy difference between the two conformations. The structure of the ribosome and Sec translocon are based on cryo-EM structures (Voorhees et al., 2014), and aside from the opening/closing of the lateral gate, are fixed in place during simulation.

Model parameterization is unchanged from previously published work (Harrington et al., 2020; Niesen et al., 2017). Beads are added to the nascent chain at a rate of 5 amino acids per second unless otherwise specified. Integration is carried out using overdamped Langevin dynamics with a timestep of 300 ns and a diffusion coefficient of 253 nm^2^/s. Translation starts 33 amino acids before the N-terminus of TM1 of the E2 protein (P663).

To measure pulling forces, translation is halted when the ribosome occupies the slip site. The dynamics of the nascent chain are then allowed to evolve for three seconds, during which time the forces exerted on the N-terminal bead are measured every 3 ms. Because the exit tunnel is truncated in the CGMD model, we add 27 amino acids to the index of the final bead to account for the unmodeled nascent chain in the omitted part of the exit tunnel. All single mutants from positions 725 to 754 are simulated. Each mutant was simulated in 100 independent runs in order to ensure the estimate for the mean pulling force has low statistical error. Data from trajectories in which TM1 does not adopt the correct Nout topology were not included in the analysis.

The ratios of topomeric isomers were obtained independently from the force-measuring simulations. To observe the final topology of TM2 in the membrane, the polyprotein was translated from amino acid P663 to I837 without pausing. Restraints were placed on TM1 to ensure it adopts the correct topology. No restraints were placed on TM2. Upon translation of the final bead, the nascent chain was released from the ribosome and simulation continued. After 3 seconds, the topology of the protein was recorded. Simulations were repeated 100 times to obtain the probability of TM2 integrating per mutant.

### Atomistic simulations of polyprotein TM2 in the translocon

An atomistic model was built of polyprotein TM2 in the Sec translocon using the cryo-EM structure with PDB code of 6ITC as an initial template (Ma et al., 2019). This structure captures a polypeptide entering a lipid bilayer through the lateral gate of SecY. The translocating peptide sequence in the structure was replaced with the polyprotein sequence using MODELLER (Webb and Sali, 2016), aligning the TM2 helix in polyprotein with the helical pro-OmpA signal sequence that is exposed to the lipid bilayer in the cryoEM structure. Specifically, residue M1 of the peptide was replaced with residue E723 of the polyprotein. Two alternative structures were generated by shifting the mapping of TM2 amino acids two amino acids forwards. Specifically, residue M1 of the peptide was replaced with either residues L725 or P727. Unresolved amino acids in the translocating peptide were modelled in, also using MODELLER. The antibodies and GFP present in the cryoEM structure were removed. The resulting structure was then solvated with TIP3P water and POPC lipids, filling a simulation box that extended 10 Å beyond the protein complex. The structure was energetically locally minimized, and then the α carbons in SecA more than 15 Å from the translocating peptide were fixed in place with 1 kcal/mol/Å^2^ harmonic restraints. Molecular dynamics simulations were then run for 150 ns with a 1.5 fs timestep at 300K and 1 atm using Desmond (Bowers et al., 2006). Root mean square fluctuations of the amino acid alpha carbons were calculated using the final 100 ns of the trajectory.

## Acknowledgments

We thank Christiane Hassel, David Frank Miller, Jun Liu, and Douglas Rusch for technical input and assistance. We also acknowledge the support of the Indiana University Flow Cytometry Core Facility and the Indiana University Center for Genomics and Bioinformatics. This research was supported in part by grant from the National Institute of Allergy and Infectious Diseases (NIAID) to J. P. S. (R21AI142383) as well as from grants from the National Institute of General Medical Sciences to T. F. M. (R01GM125063) and J. P. S. (R01GM138845). Simulations were performed using resources of the National Energy Research Scientific Computing Center (NERSC), a U.S. Department of Energy Office of Science User Facility.

## Author Contributions

J.P.S, W.D.P., S.M., and T.F.M. III designed the experiments. P.J.C., H.R.H., K.E.D., and W.D.P produced the genetic constructs and carried out the cellular and genetic experiments. C.P.K. and P.J.C. analyzed and curated the DMS data. M.H.Z. carried out and analyzed CGMD simulations. J.P.S. wrote the manuscript with editorial input from the other authors.

