## Supplemental Materials for "Coordination of -1 Programmed Ribosomal Frameshifting by Transcript and Nascent Chain Features Revealed by Deep Mutational Scanning"

*This File Includes:*

Figure S1
Figure S2
Figure S3
Figure S4
Figure S5
Figure S6

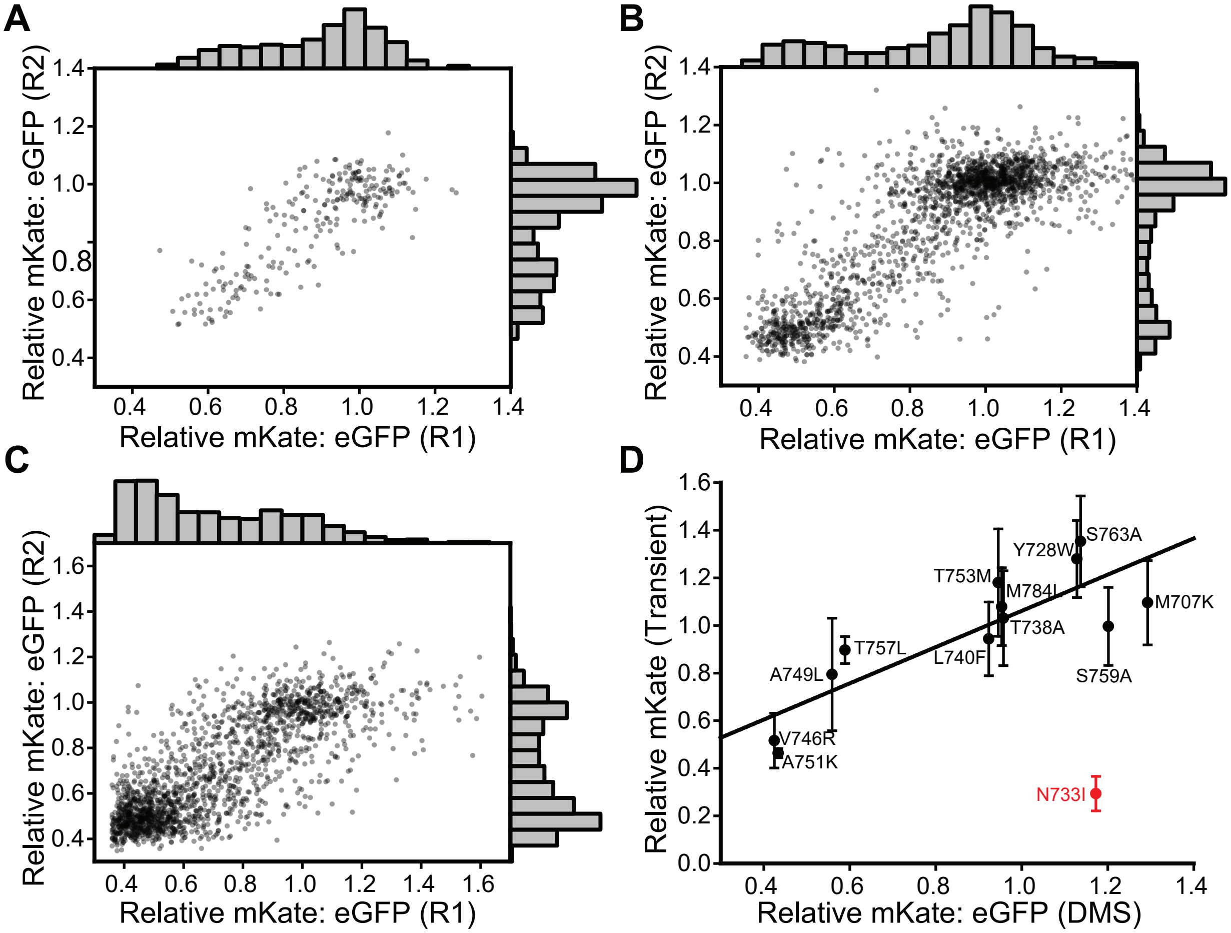


**Figure S1. Precision and accuracy of relative mKate: eGFP intensity ratios determined by deep mutational scanning.** A) For the single nucleotide variants described in Figure 2, the relative mKate: eGFP intensity ratios determined from two independent biological replicates are plotted against one another (Pearson's R = 0.83). Histograms showing the distribution of values for each replicate are shown for reference. B) For the missense mutations described in Figure 3 (redundant codons averaged), the relative mKate: eGFP intensity ratios determined from two independent biological replicates are plotted against one another (Pearson's R = 0.83). Histograms showing the distribution of values for each replicate are shown for reference. C) For codon substitutions described in Figure 6, the relative mKate: eGFP intensity ratios determined from two independent biological replicates are plotted against one another (Pearson's R = 0.79). Histograms showing the distribution of values for each replicate are shown for reference. D) A series of individual SINV structural polyprotein -1PRF reporter variants were transiently expressed in HEK293T cells, and flow cytometry was used to measure mKate intensity for each variant among positively transfected cells (eGFP+). mKate intensities are normalized relative to WT and plotted against the corresponding mKate: eGFP ratio determined by deep mutational scanning. A Grubb's outlier test identified N733I (red) as an outlier based on the residuals from a fit of the entire data set. The black line represents a linear fit of the other twelve mutants (outlier excluded, Pearson's R = 0.85).

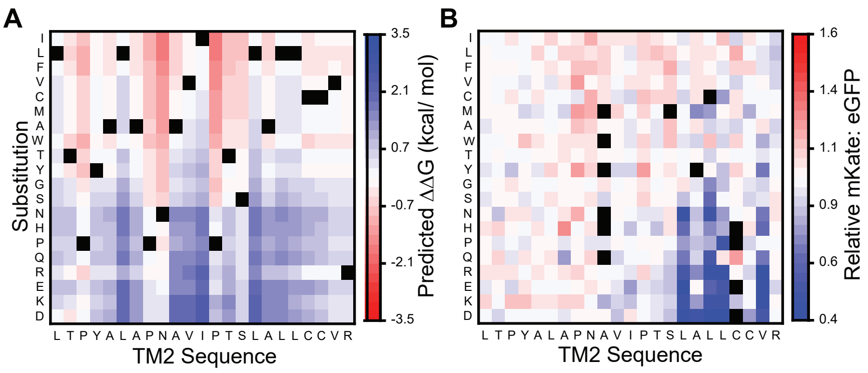


**Figure S2. Effects of missense mutations on topological energetics.** A) A Heatmap depicts the effects of each missense mutation (y-coordinate) at each position in TM2 (x-coordinate) on the predicted transfer free energy of the helix from the translocon to the membrane as determined by the ΔG predictor. B) For the sake of comparison, we also show a heatmap depicting the effects of each missense mutation (y-coordinate) at each position in TM2 (x-coordinate) on the relative mKate: eGFP intensity ratio as determined by deep mutational scanning. A value of 1.0 (white) corresponds to the value of WT. Black squares indicate a lack of data.

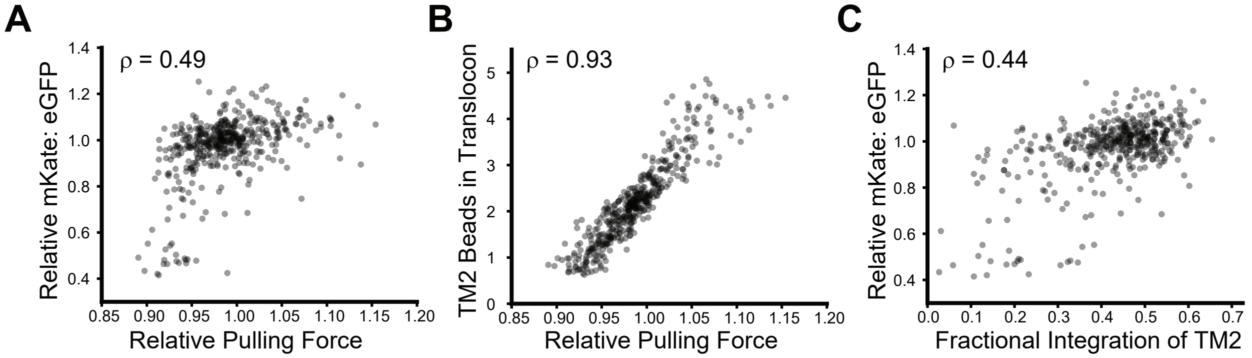


**Figure S3. Relationship between pulling force, membrane integration, and frameshifting.** The effects of mutations on frameshifting are plotted in relation to their effects on the topological properties and pulling forces on the nascnet chain in CGMD simulations. Each point represents one of the 454 possible missense mutations in TM2. A) A plot of relative mKate: eGFP intensity ratio measurements against the average pulling force values normalized relative to WT shows that mutations that increase pulling forces generally increase frameshifting, though the realtionship is potentially non-linear. B) The average number of TM2 beads within the translocon when the ribosome occupies the slip-site is plotted against the average pulling force normalized relative to WT. The apparent linear relationship suggests the exposure of the nascent chain to the protein conducting channel and/ or the membrane core are involved in the generation of pulling force. C) Relative mKate:eGFP intensity ratio measurments are plotted against the fraction of TM2 that ultimately undergoes membrane integration following translation of the slip-site. The modest correlation shows that mutations that increase the membrane integration of TM2 generally increase frameshifting. The spearman's correlation coefficient (ρ) is shown for each data set.

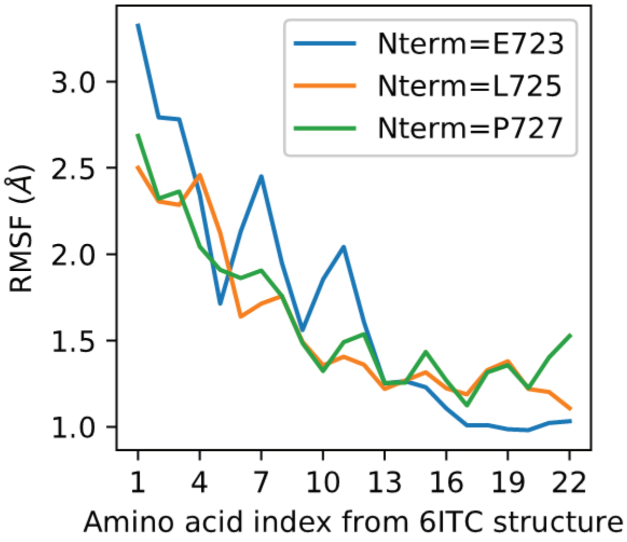


**Figure S4. Conformational dynamics of various atomistic models of TM2 within the translocon.** The sequence of the polyprotein was mapped onto the nascent chain within a cryo-EM structure of a translocation intermediate (PDB ID 6ITC) begining at three different residues (E723, L725, and P727) in order to place TM2 near its approximate position during frameshifting. These maps were used to generate three atomistic models of the nascent chain that were each relaxed for 150 ns in the context of the translocon and an explicit lipid bilayer (see *Methods*). All three models exhibit similar topological properties and display the same trend of decreasing RMSF from the N- to C-terminal residues of TM2. These results show that the observed structural and dynamic properties of this translocation intermediate are unlikely to be sensitive to subtle variations in its positon with respect to the translocon.

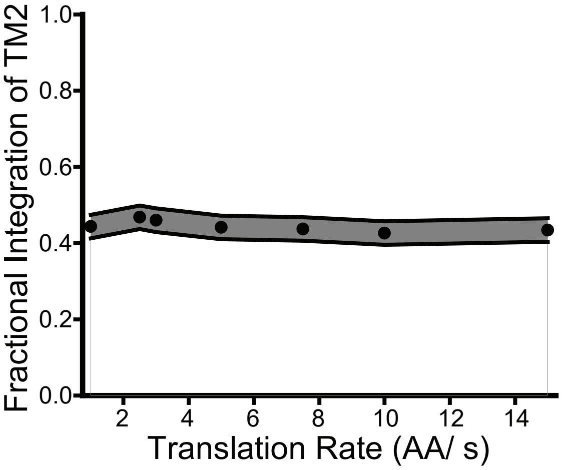


**Figure S5. Impact of translation kinetics on the translocon-mediated membrane integration of TM2.** Coarse-grained molecular dynamics simulations of the translocon-mediated cotranslational folding of the SINV structural polyprotein (300 trajectories per condition) were carried out with a varying rate of translation. The fraction of trajectories in which TM2 spans the membrane following translation of the slip-site is plotted against the translation rate. The gray bar depicts the bounds of the 90% confidence interval. These results suggest that translation kinetics have minimal impact on the final membrane integration efficiency of TM2.

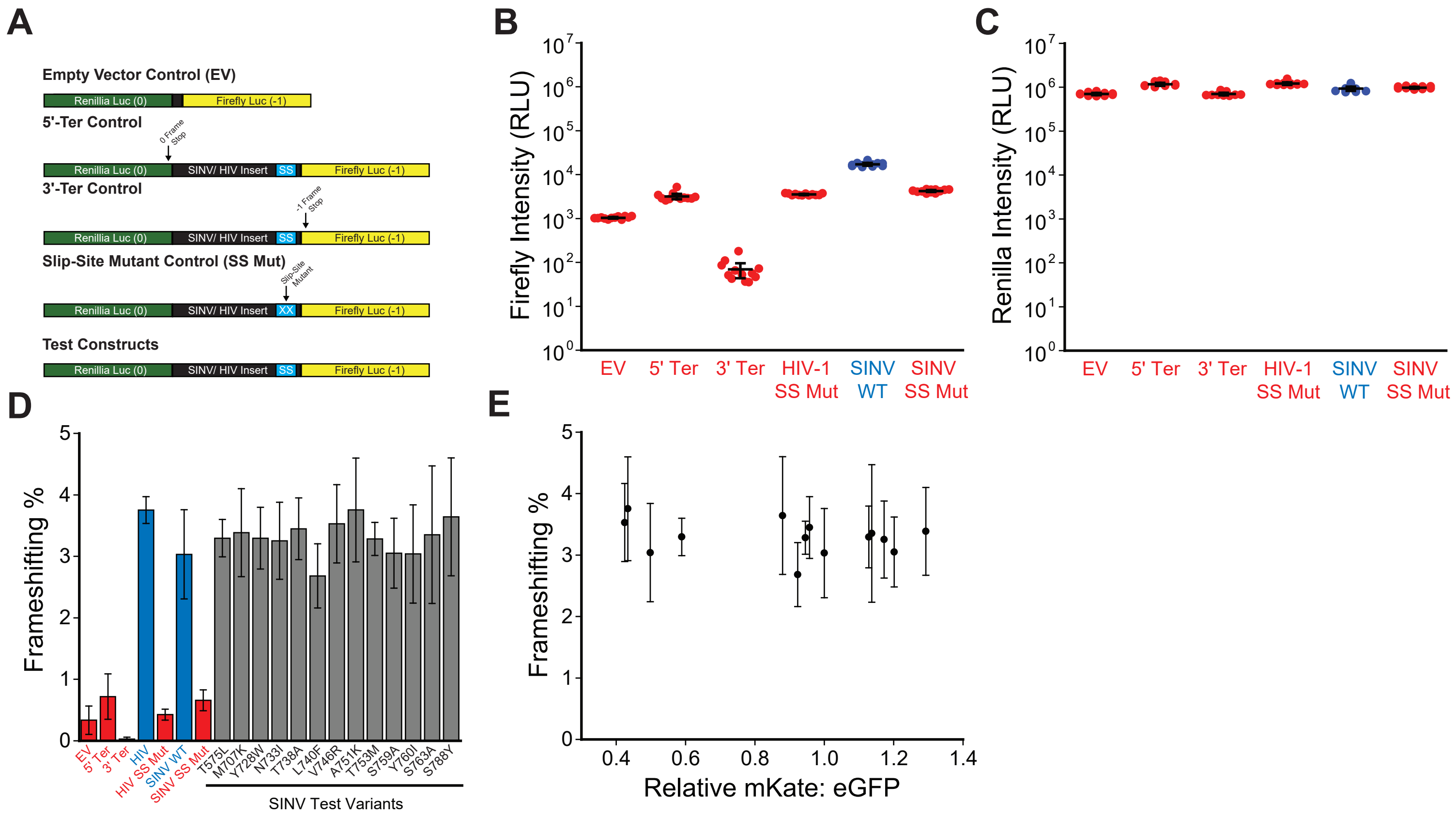


**Figure S6. Impact of Mutations on -1PRF Monitored by a Dual-Luciferase Reporter.** A) Schematic diagrams depict the relative orientations and reading frames (0 or -1) of renilla luciferase (green) and firefly luciferase (yellow) cassettes in relation to the insert (150 bp HIV-1 gag-pol motif or the 1.7 kb SINV fragment) for both negative controls and test constructs. Inserts were flanked by self-cleaving P2A linkers and cryptic splice sites within the renilla luciferase cassette were removed as was described in Khan et al., 2019. B) Replicate firefly luciferase intensity measurements from select control reactions are shown from a single representative biological trial (12 readings from 4 independent transfections each). C) Replicate renilla luciferase intensity measurements from select control reactions are shown from a single representative biological trial (12 readings from 4 independent transfections each). D) A bar graph shows the average -1PRF efficiency for each positive control (blue), negative control (red), and SINV test variant (gray) from three biological replicates in HEK293 cells. Error bars reflect the standard deviation. Efficiency values were calculated by dividing the ratio of the firefly to renilla luciferase intensities for each construct by that of a no-insert-control containing both luciferase cassettes in the 0-reading frame, as was previously described (Khan et al., 2019). E) Average frameshifting effiicency measurments along with stadard deviation values for SINV test variants shown in panel D are plotted against their corresponding relative mKate: eGFP values as was determined by deep mutational scanning (DMS). Though fluorescence reporters suggest these mutations impact the efficiency of -1PRF (Figures 3, & S1), dual luciferase measurments are relatively insensitive to their effects. Unlike the fluorescence reporters used for DMS, dual-luciferase reporters bear a reporter domain upstream of the region encoding the E3 signal peptide. Based on this consideration, we suspect the disagreement between these assays arises as a result of the inefficient ER-targetting of the SINV structural polyprotein insert in the context of dual-luciferase reporters.
